# Hsp90 buffers behavioral plasticity by regulating *Pdf* transcription in clock neurons of *Drosophila melanogaster*

**DOI:** 10.1101/2025.03.26.645458

**Authors:** Angelica Coculla, Carlina Feldmann, Maite Ogueta, Sina Mews, Roland Langrock, Ralf Stanewsky

## Abstract

Circadian rhythms are prevalent on Earth and temporally organize behaviour and physiology of organisms to occur in species-specific ‘temporal niches’. However, species differ in how strictly individuals are controlled by their circadian clock, suggesting that it may offer a selective advantage for an individual to extend its temporal niche under certain circumstances, for example during stressful environmental conditions. A potential mechanism controlling temporal niche adherence involves the evolutionary capacitor and chaperon protein HSP90, known to assist the proper folding of important signalling molecules. If HSP90 becomes rate limiting (e.g., under environmental stress) hidden genetic variation will be expressed, producing novel and potentially beneficial phenotypes for the individual. While this role of HSP90 is well established for morphological traits, we show here that it extends to regulation of temporal behavioural patterns. We show that within a small subset of clock neurons in the fly brain, HSP83, the fly homologue of HSP90, mitigates inter-individual behavioural plasticity. We provide evidence for the requirement of HSP83 for efficient transcription of the gene encoding the circadian neuropeptide <u>P</u>igment <u>D</u>ispersing <u>F</u>actor (PDF), and for correct PDF accumulation in central clock neurons. Strikingly, *Hsp83* mutants affect synchronized oscillations of the clock protein PERIOD (PER) in subsets of circadian clock neurons in the same way as flies without PDF, further supporting a role of *Hsp83* in regulating *Pdf*. Our findings therefore provide a mechanistic explanation for HSP83 function in regulation of behavioural plasticity, and offer an explanation for how to restrict temporal niche extension to stressful environmental conditions.

## Introduction

Animals restrict their behavioral activities to specific times of the day or night, and these behaviours are regulated by self-sustained circadian clocks. Environmental rhythms, like the daily light:dark and concomitant temperature changes synchronize the circadian clock with the external time. This allows animals to exert behaviours at optimal times of day, often by anticipating the actual environmental change. Circadian clocks therefore contribute to overall fitness, for example through minimizing energy expenditure, increasing mating probability, or avoiding predators (van der Vinne et al., 2019). In *Drosophila melanogaster*, daily activity rhythms are orchestrated by a network of ∼240 clock neurons in the fly brain (Reinhard et al., 2024). Within these neurons transcriptional and translational molecular feedback loops of key circadian clock genes generate ∼ 24 h molecular oscillations, which are translated into neural activity rhythms, rhythmic neuropeptide release, as well as into rhythmic activation of clock-controlled output genes, ultimately regulating rhythmic behaviour and determining the crepuscular temporal niche of the fruit fly (Helfrich-Förster and Reinhard, 2025; Tataroglu and Emery, 2015). The core feedback loop is initiated by a heterodimer formed of the two transcription factors CLOCK (CLK) and CYCLE (CYC), which binds to E-box elements upstream of the *period* (*per*) and *timeless* (*tim*) genes, thereby activating their transcription. PER and TIM proteins gradually accumulate in the cytoplasm and eventually enter the nucleus to repress CLK-CYC activity and *per/tim* transcription. After degradation of PER and TIM, CLK-CYC can reinitiate *per* and *tim* transcription and the feedback loop repeats itself every 24 h. CLK and CYC are also part of an interlocked feedback loop, in which they activate the transcription of the genes encoding the basic-zipper transcription factors VRILLE (VRI) and PAR DOMAIN PROTEIN 1ε (PDP1ε), which repress, or activate *Clk* transcription, respectively (Tataroglu and Emery, 2015). Inactivating *vri* or *Pdp1ε* in the clock neurons abolishes locomotor activity rhythms without interfering with molecular clock function, demonstrating that the interlocked feedback loop is mainly responsible for clock output (Benito et al., 2007; Gunawardhana and Hardin, 2017; Zheng et al., 2009). The neuropeptide pigment dispersing factor (PDF) is expressed in a subset of the clock neurons, four small as well as four large ventral lateral neurons (s-LNv and l-LNv) and required for maintaining behavioural rhythmicity in constant conditions (Renn et al., 1999). While PDF is not required for the maintenance of molecular oscillations in the clock neurons, it does coordinate the phase and amplitude between different neuronal groups, which is essential for maintaining rhythmic behaviour (Lin et al., 2004; Yoshii et al., 2009). In turn, PDF expression is posttranscriptionally regulated by the circadian clock via VRI, which is required for normal accumulation of *Pdf* RNA and peptide levels. As a consequence, the daily plasticity rhythms of the s-LNv terminals in the dorsal brain are abolished, most likely contributing to the behavioural arrhythmicity of *vri* mutant flies (Gunawardhana and Hardin, 2017).

Interestingly, despite the remarkable conservation of circadian clock genes and mechanisms in the animal kingdom, the degree of rhythmically behaving individuals within a certain species is surprisingly variable. While in *D. melanogaster* nearly each individual displays robust behavioural rhythms, we have recently shown that in the flour beetle *Tribolium castaneum* only 30-50 % of the individuals behave rhythmically (R et al., 2024). This difference between species may allow for adjustments of circadian clocks to the specific ecological demands of a particular species. For example, under environmental stress (e.g., drought, excessive heat), it could be advantageous not to be restricted to a narrow temporal niche, but to extend it, or even to be active at irregular times. In a population, these individuals would have a fitness advantage under certain environmental conditions, an evolutionary strategy termed as bet hedging (Cohen, 1966). On the other hand, even a largely homogenously rhythmic species like *D. melanogaster* could possess buffered standing genetic variation. Such cryptic genetic variation (CGV) would only emerge under stressful environmental conditions, leading to inter-individual behavioural plasticity. For morphological traits, the evolutionary capacitor HSP90 has been implicated in the release of such CGV, for example in fruit flies and cave fish (Rohner et al., 2013; Rutherford and Lindquist, 1998). HSP90 is a chaperon protein assisting in the folding of important signalling molecules, such as transcription factors, kinases, and ubiquitin ligases. Under non-stressful conditions, HSP90 is not rate-limiting, allowing correct protein folding, even in the presence of mild hypomorphic missense mutations. Protein folding is sensitive to environmental stress and HSP90 becomes rate limiting under these conditions, thereby unmasking such cryptic genetic polymorphisms, with potential beneficial phenotypes for the individual (Rohner et al., 2013; Rutherford and Lindquist, 1998). There is also evidence that HSP83 acts as chaperon for epigenetic factors, notably proteins encoded by the *trithorax* group (TrxG) genes, thought to be involved in the maintenance of active chromatin states (Paro et al., 1998; Sollars et al., 2003). It was shown that HSP90 depletion also resulted in morphological plasticity in isogenic *Drosophila melanogaster* and *Arabidosis thaliana* plants together suggesting an epigenetic mechanism for HSP90 capacitor function (Sollars et al., 2003). In contrast to morphological traits, it is questionable whether release of CGV or misfolding of epigenetic factors by limited availability of HSP90, also contribute to behavioural plasticity. An increase of behavioural variability under stressful environmental conditions by either mechanism would allow certain animals of a population to extend their temporal niche, potentially offering a selective advantage for these individuals. A previous study conducted in *Drosophila* pointed to a potential role of HSP90 (HSP83 in flies) as capacitor of behavioural plasticity in circadian locomotor activity (Hung et al., 2009). Reduction of HSP83 resulted in a marked increase of inter-individual behavioural plasticity without apparent alterations of the underlying molecular clock oscillations (Hung et al., 2009). Here, we aimed to identify the cellular substrates of circadian HSP83 function. Using an inducible *HSP83* CRISPR allele, we show that behavioural variability increases when *Hsp83* function is restricted to clock neurons only. *Hsp83* is required for normal *Pdf* transcription and accumulation of PDF in the s-LNv cell bodies and projections. Strikingly, *Hsp83* mutation resembles the effect of *Pdf^01^* loss-of-function mutants on PER expression rhythms in subsets of PDF-negative neurons indicating faulty synchronisation between different clock neuronal groups. Our results show that HSP83 is required for normal PDF signalling within the circadian clock and suggests that this chaperon is important for the proper folding of proteins that regulate PDF. Interestingly, it has recently been shown that DNA sequence differences in the non-coding, upstream region of the *Pdf* gene between two Drosophila species (*D. sechellia* and *D. melanogaster*) translate into different PDF peptide levels and distinct behavioural patterns (Shahandeh et al., 2024). Therefore, transcriptional regulation of a single circadian neuropeptide gene (*Pdf*), can explain behavioural circadian plasticity between different *species* and between *individuals* of a population.

## Results

### *Hsp83* mutants exhibit inter-individual behavioral variability

To confirm that interference with *Hsp83* function indeed increases behavioral variability of circadian behaviors, we analyzed locomotor activity of *Hsp83* alleles previously reported to affect behavior (Hung et al., 2009). We used the same three loss-of-function alleles (*Hsp83^e6A^*, *Hsp83^e6D^*, and *Hsp83^j5c2^*) and one hypomorphic allele (*Hsp83^08445^*). As expected, all loss-of-function alleles were homozygous lethal, and no transheterozygous offspring was obtained after crossing them to a deficiency removing the entire *Hsp83* locus (Tab S1). In contrast *Hsp83^08445^* flies are homozygous viable and generate viable transheterozygous offspring after crossing to the deficiency and the loss-of-function alleles ((Hung et al., 2009), Tab S1).

We tested the locomotor behavior of the homozygous and transheterozygous *Hsp83* mutants and observed significant changes of the rhythmic locomotor activity patterns in the mutants compared to the controls (wild type flies and *Hsp83^mut^/*+) both in LD and DD (Fig 1A). While control flies showed similar patterns in locomotor activity during entrainment and free running condition, the homozygous and trans-heterozygous offspring showed significant intra- and inter-individual differences (Fig 1A). At the population level, mutants have less well-defined activity peaks compared to their respective controls in both LD and in DD (Fig 1B, C).

**Fig 1:**
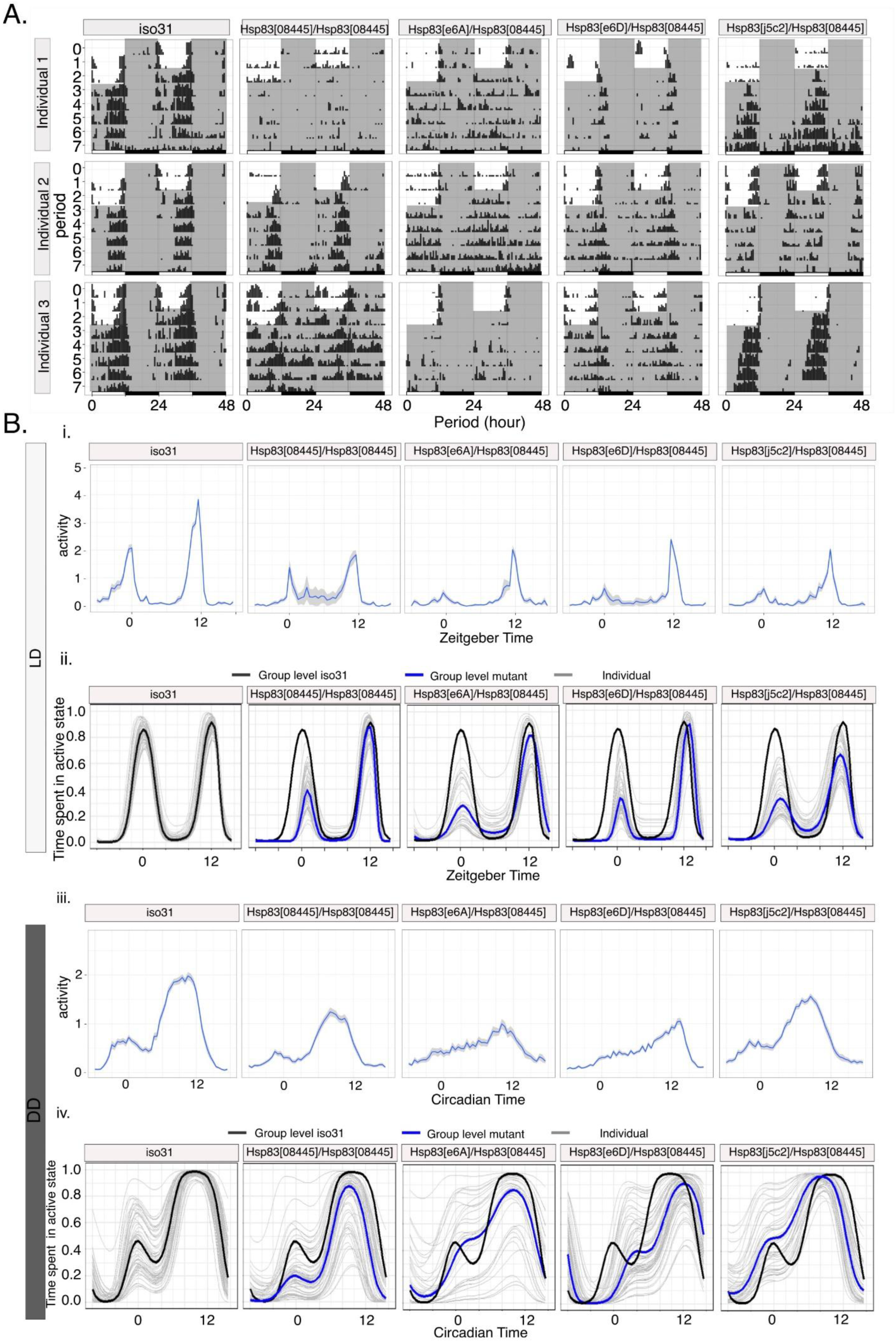
Reducing Hsp83 function increases behavioral variability in *Drosophila melanogaster*. A) Representative actograms of *Hsp83* mutant flies compared to iso31. White areas represent ‘lights on’ and gray areas ‘lights-off’. B) Population Activity (top) and HMM-implied percentage of time spent in the active state (bottom) of *Hsp83* mutants and their controls in LD. C) Population Activity (top) and HMM-implied percentage of time spent in the active state (bottom) of Hsp83 mutants and their controls in DD. See also Tab 1, 2, S2.

### *Hsp83* mutants impair synchronization during light:dark cycles

Inspection of the average activity plots showed that *Hsp83* mutants display an amplitude reduction of the morning (M) and evening (E) peaks, indicating an impairment of synchronization to LD cycles (Fig 1B). To quantify this behavior, we calculated an Entrainment index (EI), reflecting the ratio of M and E peak activity over the activity during the entire 24 hours ((Gentile et al., 2013), Materials and Methods). Compared to the wild type controls, M and E peak synchronization was reduced in all *Hsp83* mutant combinations, particularly around lights on (Tab 1). Moreover, in two heterozygous loss-of-function mutants (*Hsp83^e6A^/+,* and *Hsp83^e6D^/+*) morning synchronization was reduced, which most likely is attributable to the generally higher M-peak variability, and indicating a dominant effect of *Hsp83* depletion (Tab 1). To investigate inter-individual variability more comprehensively, we analyzed LD and DD behavior separately using hidden Markov models (HMMs) as an alternative approach. An HMM is a statistical model that allows to identify not directly observable states (i.e., hidden states) based on the observable data, in our case locomotor activity counts (Feldmann et al., 2023). These models are frequently applied in animal ecology to infer unobserved behavioral processes (e.g., active and inactive) from observed time series data (e.g., tracking data or locomotor activity) (Bauhus et al., 2024; Feldmann et al., 2023; Nagel et al., 2021), and have been previously used with activity data in Drosophila to identify sleep stages (Wiggin et al., 2020). Here, we used HMMs to distinguish two states: active and inactive (see Materials and Methods for details). Comparing average histogram plots with HMM analysis, revealed the latter accurately reports the crepuscular behaviour of wild type flies displayed during LD cycles (Fig 1B, compare top and bottom graphs) (Feldmann et al., 2023). Focusing on the model-implied percentage of time spent in the active state shortly before and after each environmental transition, we observed that overall *Hsp83* mutants were more likely to be variable compared to the wild type (Fig 1B, Tab 1). Interestingly, the time spent in the active state was more variable within the population also in the heterozygous mutant flies, with the exception of *Hsp83^08445^/+* and *Hsp83^e6A^/+* (Tab 1). In summary, the HMM analysis revealed that inter-individual behavioral plasticity around the lights-on and lights-off transitions is increased in all *Hsp83* mutants analyzed here. Moreover, at the population level this variability results in drastically reduced M-peak activity in *Hsp83* mutants, suggesting that most of the mutant flies are not in the active state around lights-on (Fig 1B, Tab 1).

**Tab 1.**
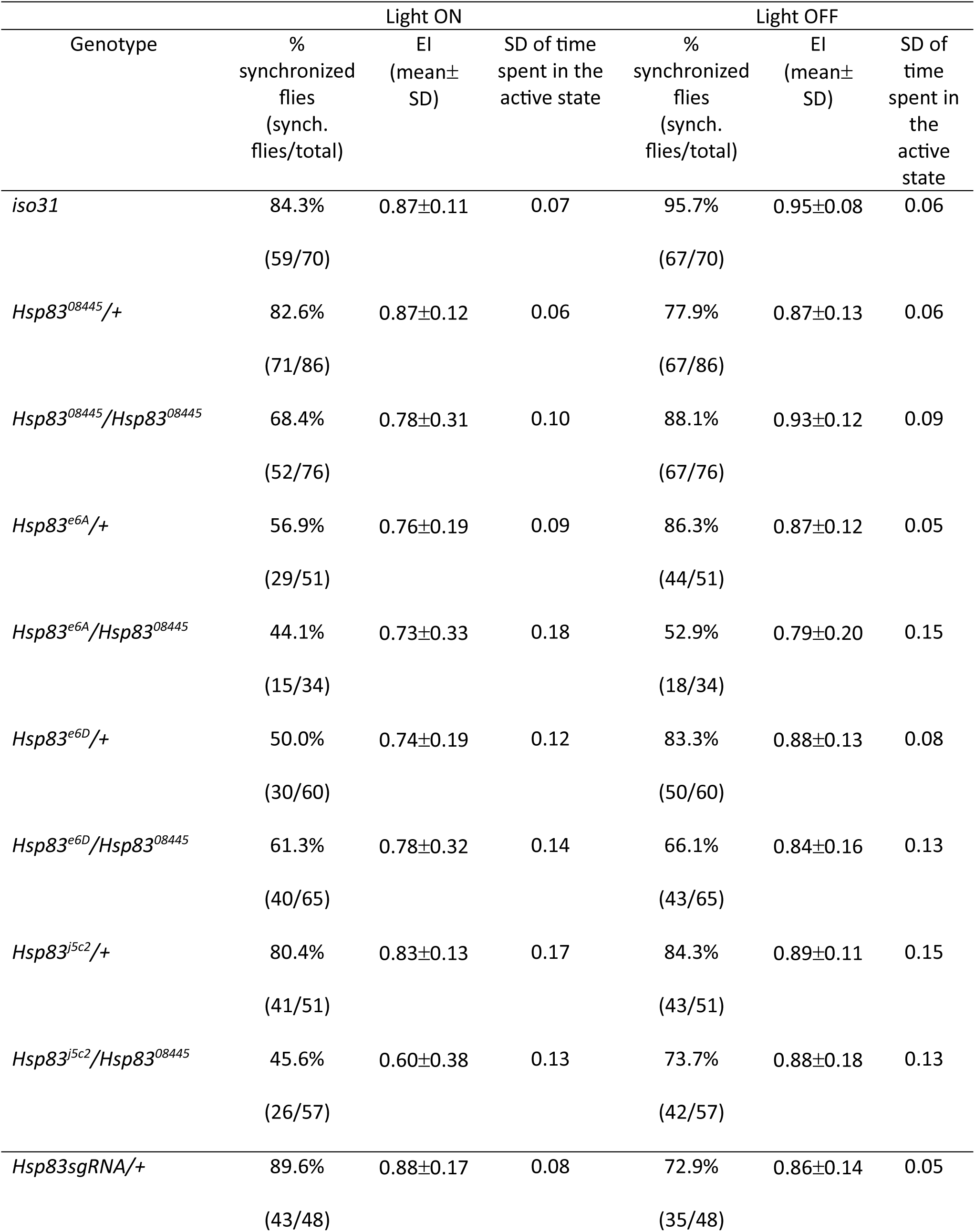
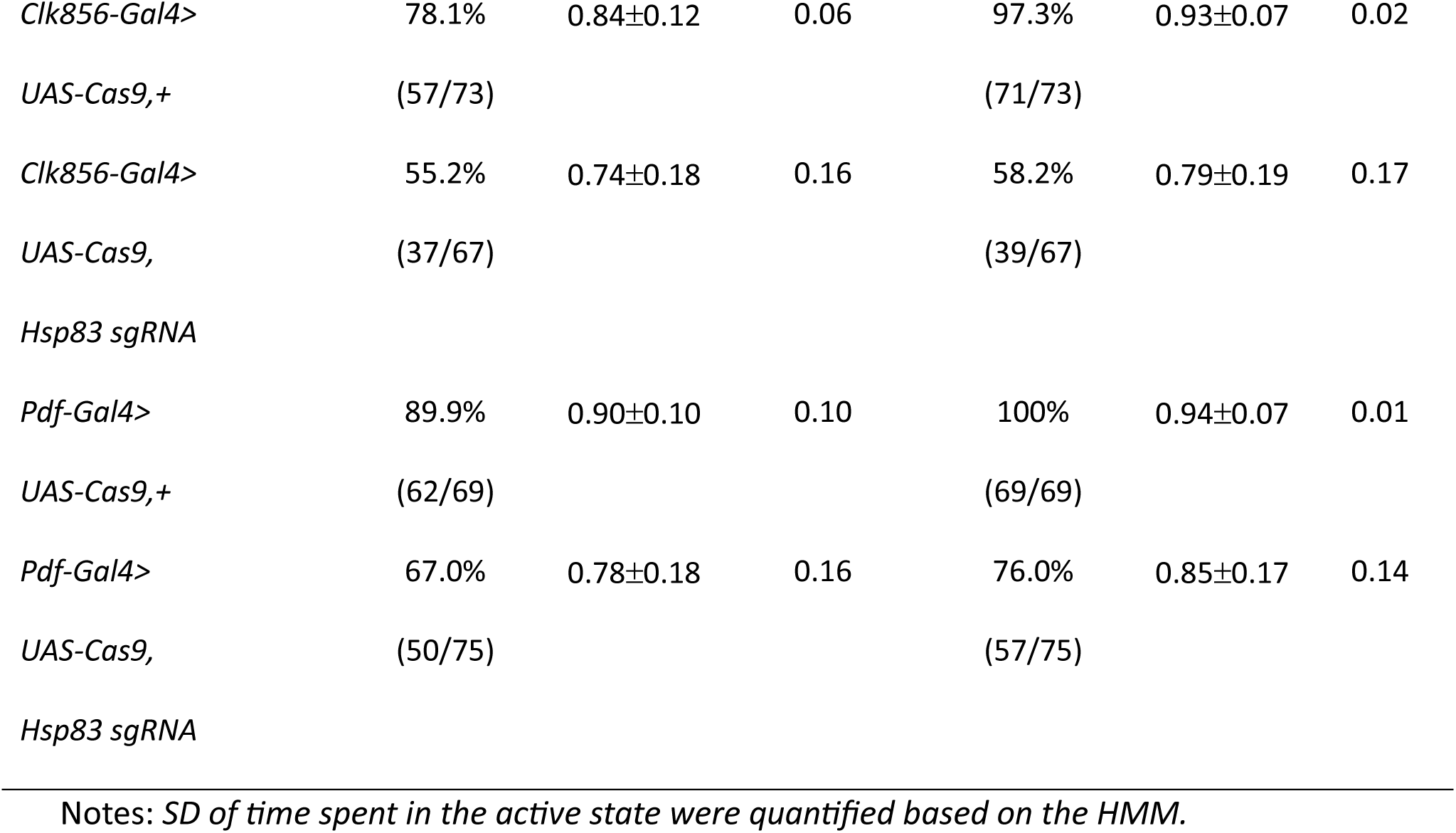
Analysis of locomotor activity during LD.

### *Hsp83* mutants increase within-group behavioral variability in constant darkness

Inspection of individual actograms also revealed variable activity patterns between *Hsp83* mutant individuals in DD (Fig 1A). To quantify this variability, we first analyzed rhythmicity and period-length, because a previous study reported a significant reduction of rhythmicity in *Hsp83* mutants in DD (Hung et al., 2009). In our hands, *Hsp83* mutants did not show a reduction in overall rhythmicity compared to the controls, except for *hsp83^e6A^/hsp83^08445^* flies, which exhibited a ∼20% reduction and reduced rhythmic strength (Tab 2). Moreover, as previously reported, all *Hsp83* mutant combinations showed normal free-running period lengths (Tab 2, (Hung et al., 2009)), indicating that core clock function is not affected. Although overall rhythmicity appears not, or only mildly affected, the individual and group activity plots indicate substantial inter-individual plasticity of the DD activity pattern in *Hsp83* mutants (Fig 1A, C). To determine if variability under constant conditions is indeed increased in *Hsp83* mutants, we quantified the activity phase of each individual on two consecutive days. For wild type flies, activity peaks on consecutive days will occur around the same time, leading to a high degree of ‘phase coherence’ within the population. Based on the sometimes erratic behavior observed in individual actograms of *Hsp83* mutants (Fig 1A), we expected to see a reduction in phase coherence in the mutants. Indeed, inspection of the circular phase plots revealed that all *Hsp83* mutant combinations showed reduced phase coherence (length of the colored vector emerging from the center of each circle) (Fig. 2; Tab S2). Increased inter-individual variability was independently supported by the HMM analysis, because *Hsp83* mutants exhibit increased variability of the model-implied percentage of time spent in the active state around the subjective evening peak (Fig 1C, Tab S2). Combined, the results show that *Hsp83* mutant individuals display most of their activity at different times during the 24-hr day.

**Tab 2:** Analysis of locomotor activity in DD.

**Tab 2: Analysis of locomotor activity in DD.**
| Genotype | % Rhythmic<br>(n of rhythmic<br>/total flies) | Period (h) ±<br>SEM | RS ± SEM |
| --- | --- | --- | --- |
| *iso31* | 98.6% (69/70) | 23.7 ± 0.03 | 4.1 ± 0.08 |
| *Hsp83<sup>08445</sup>/+* | 98.8% (85/86) | 23.6 ± 0.02 | 4.1 ± 0.10 |
| *Hsp83<sup>08445</sup>/Hsp83<sup>08445</sup>* | 97.4% (74/76) | 23.5 ± 0.13 | 3.4 ± 0.13 |
| *Hsp83<sup>e6A</sup>/+* | 100 % (51/51) | 23.9 ± 0.06 | 4.2 ± 0.13 |
| *Hsp83<sup>e6A</sup>/Hsp83<sup>08445</sup>* | 73.5% (25/34) | 23.7 ± 0.10 | 2.9 ± 0.21 |
| *Hsp83<sup>e6D</sup>/+* | 98.3% (59/60) | 24.3 ± 0.03 | 3.9 ± 0.07 |
| *Hsp83<sup>e6D</sup>/Hsp83<sup>08445</sup>* | 98.5% (64/65) | 24.2 ± 0.05 | 3.3 ± 0.14 |
| *Hsp83<sup>j5c2</sup>/+* | 98.0% (50/51) | 23.7 ± 0.04 | 4.0 ± 0.11 |
| *Hsp83<sup>j5c2</sup>/Hsp83<sup>08445</sup>* | 96.4% (54/56) | 23.5 ± 0.07 | 4.1 ± 0.11 |
| *Hsp83 sgRNA/+* | 100 % (99/99) | 23.6 ± 0.02 | 4.5 ± 0.09 |
| *Clk856-Gal4>UAS-Cas9,+* | 98.6% (72/73) | 23.6 ± 0.03 | 4.5 ± 0.11 |
| *Clk856-Gal4>UAS-Cas9,Hsp83 sgRNA* | 50.7% (34/67) | 24.6 ± 0.35 | 2.6 ± 0.20 |
| *Pdf-Gal4>UAS-Cas9,+* | 100 % (84/84) | 23.9 ± 0.02 | 5.1 ± 0.10 |
| *Pdf-Gal4>UAS-Cas9,Hsp83 sgRNA* | 94.7% (71/75) | 23.8 ± 0.08 | 3.9 ± 0.15 |

**Fig 2:**
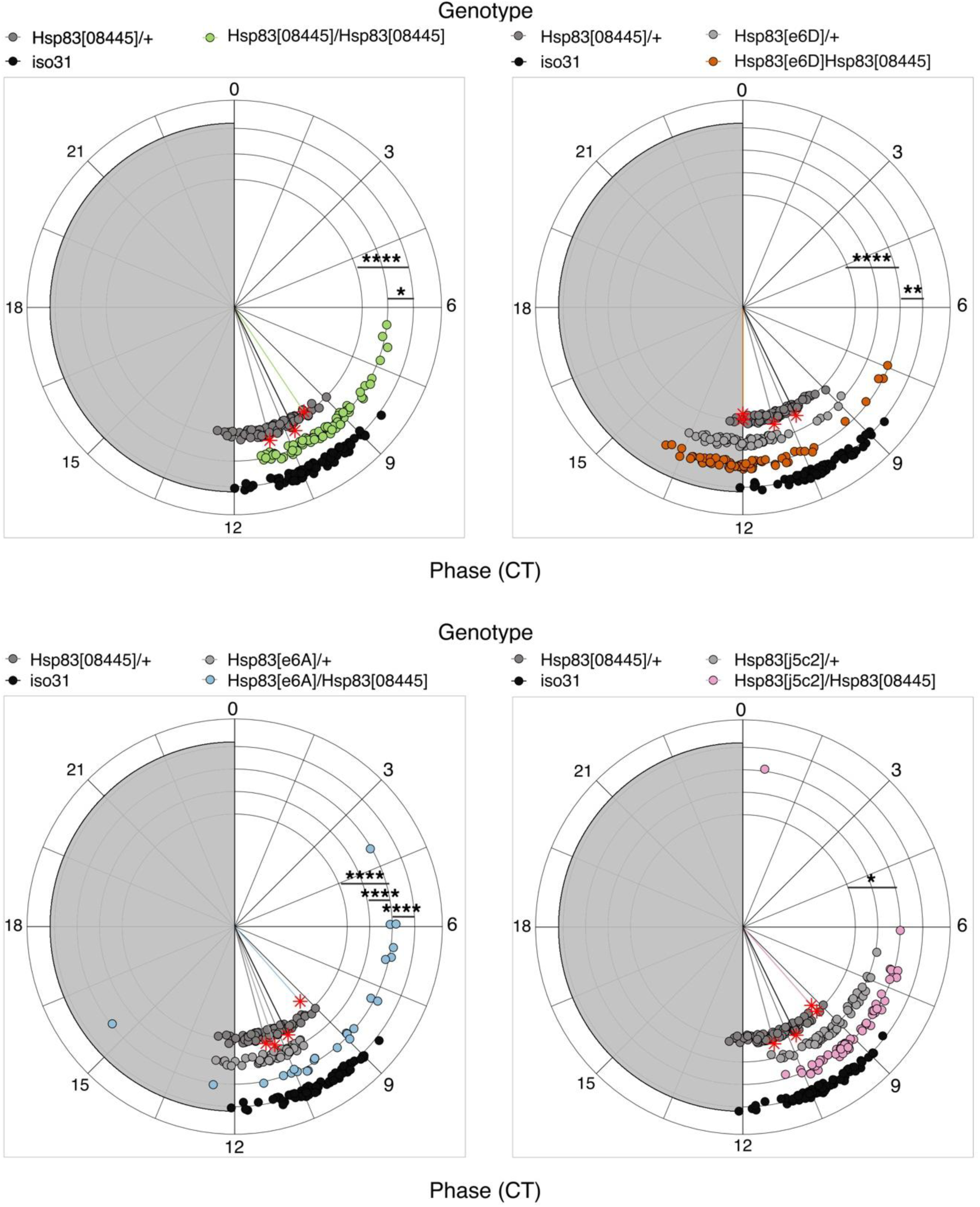
*Hsp83* mutants decrease phase coherence between individuals in constant darkness. Circular phase plots showing the peak phase in DD, calculated using the second (DD2) and the third day (DD3) in constant darkness. Each *Hsp83* mutant allele was compared with the corresponding parental control and with wild type (*iso31*). For all transheterozygous combinations, homozygous *Hsp83^08445^* females were used as mothers (including *Hsp83^08445^*/+). Colored lines represent the median phase of each group, while the length of the vector represents the standard deviation of the corresponding group (i.e., the phase coherence) (Tab S2) (Levene’s Test: F_(8,519)_= 7.92, *p*<.0001; Bartlett Test: *χ^2^*= 135.11, df=8, *p*<.0001). Shapiro Normality test, F-test and Mood test (Tab S4, S5). Note: * p < 0.05, ** p < 0.01, *** p < 0.001, **** p < 0.0001.

### *Hsp83* knock out in clock neurons increases behavioral variability

The PDF neuropeptide influences synchronization between different clock neuronal groups (Lin et al., 2004; Yoshii et al., 2009), raising the possibility that *Hsp83* depletion affects PDF, which in turn would result in desynchronized behavior. We therefore spatially restricted the disruption of *Hsp83* function, by expressing a ubiquitously expressed *Hsp83-sgRNA* line together with *UAS-Cas9* in clock neurons, thereby generating clock-neuron-specific *Hsp83* mutations (Port and Bullock, 2016). Using the *timeless-Gal4* driver, which is expressed in all peripheral clock cells in addition to clock neurons in the brain, resulted in no viable offspring (Tab S1). This indicates that the CRISPR/Cas9 meditated knock-out effectively disrupts *Hsp83* gene function, resulting in lethality as in the loss-of-function mutants described above (Tab S1). Next, we restricted *Hsp83* knock-out to all (*Clk856-gal4; UAS-Cas9*), or to the PDF-expressing subset of clock neurons (*Pdf-Gal4; UAS-Cas9*) (Fig 3A) (Gummadova et al., 2009; Renn et al., 1999), resulting in viable offspring in both cases, as expected (Tab S1).

Clock-neuron specific *Hsp83* knock-out resulted in aberrant and variable temporal locomotor activity patterns in LD and DD conditions, similar as described for the *Hsp83* mutants above (Fig 3B-D). EI quantification and HMM analysis of the behavior during LD revealed that, synchronization of the M- and E-peaks was impaired, both after *Hsp83* knock-out in all, or the PDF-clock neurons only (Fig 3C, Tab 1). Interestingly, in DD *Hsp83* knock-out in all clock neurons reduced the rhythmicity by ∼50% compared to control flies, while there was only a very subtle effect (∼6% reduction) after *Hsp83* knock-out in the PDF neurons (Tab 2). As observed for the *Hsp83* mutants, CRISPR-induced knockout in all, or the PDF neurons only, had no effect on period length (Tab 2). Next, we analyzed if *Hsp83* knock-out also affected phase coherence in DD, as indicated by the variable behavior in individual actograms (Fig 3). Indeed, circular phase and HMM analysis revealed that *Hsp83* knock-out in all, or only in the PDF subset of clock neurons significantly decreased phase coherence, demonstrating an increase of inter-individual behavioral variability (Fig. 3D, 4, Tab S2).

**Fig 3:**
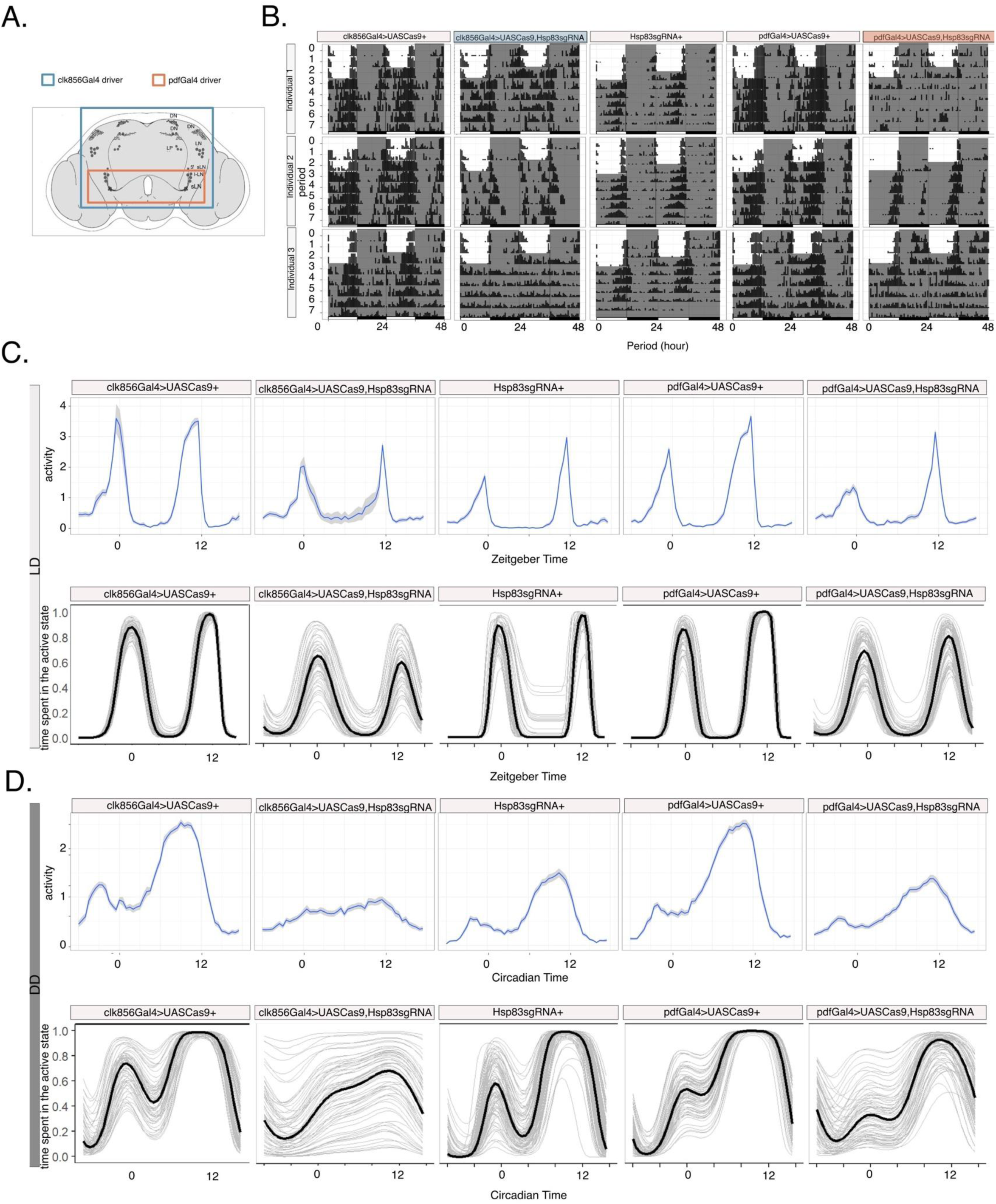
*Hsp83* knock-out in clock neurons increases behavioral variability. A) carton of the adult clock neurons in the fly brain and neurons expressing the indicated Gal4 drivers. B) Representative actograms of *Hsp83* knock-out and control flies. White areas indicate times when the lights were on, and dark areas, times when the lights were off, respectively. C) Population Activity (top) and HMM-implied percentage of time spent in the active state (bottom) of flies with *Hsp83* knock-out in all clock neurons (*Clk856>Cas9, Hsp83-sgRNAi)*, or the PDF expressing subset only (*Pdf>Cas9, Hsp83-sgRNAi)* and their controls LD (see also Tab 1). D) Population Activity (top) and HMM-implied percentage of time spent in the active state (bottom) of flies with *Hsp83* knock-out in all clock neurons (*Clk856>Cas9, Hsp83-sgRNAi)* or the PDF expressing subset only (*Pdf>Cas9, Hsp83-sgRNAi)* and their controls in DD (see also Tab S2).

**Fig 4:**
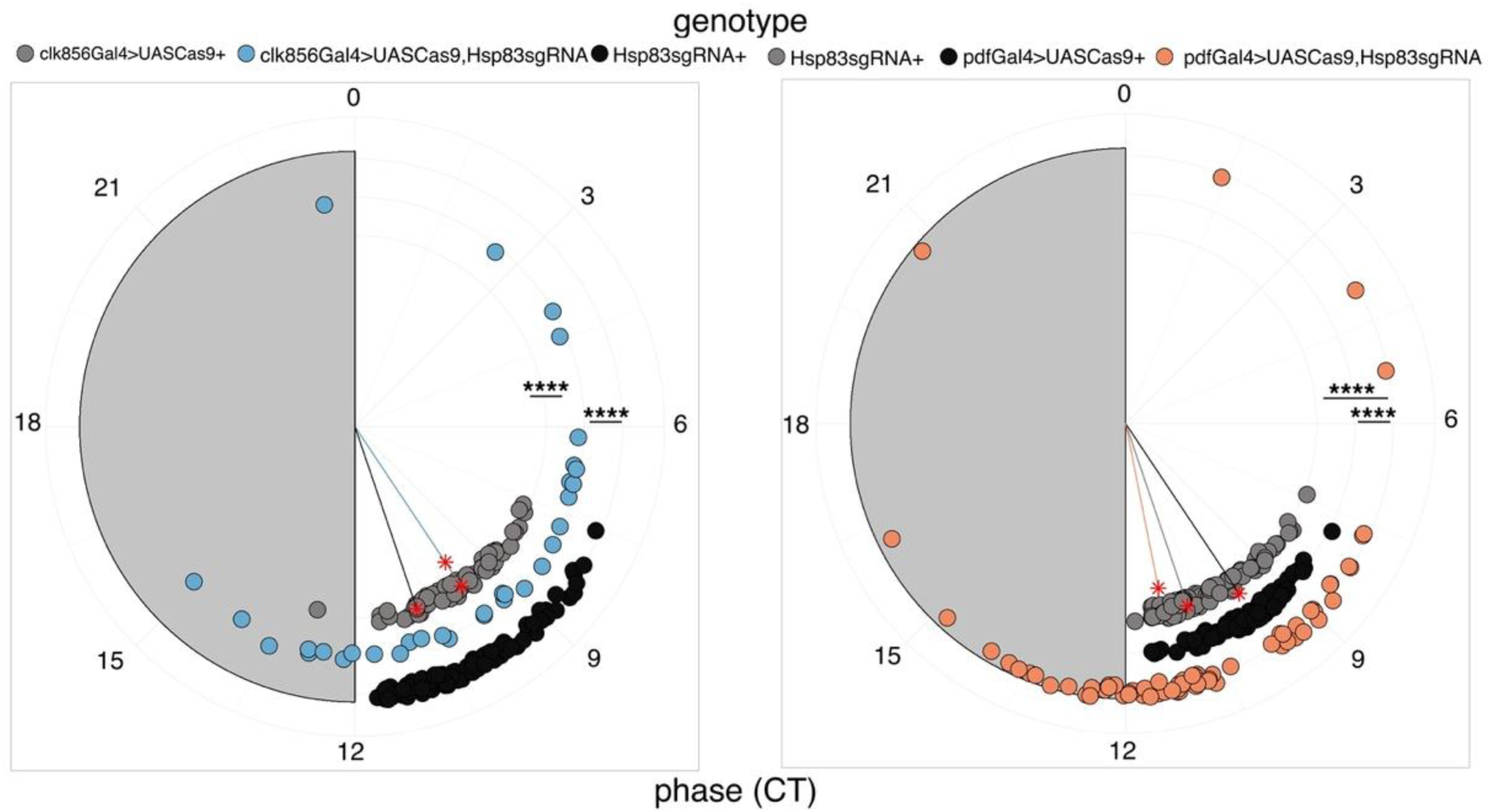
*Hsp83* knock-out in clock neurons decreases phase coherence between individuals. Circular phase plots showing the peak phase in DD for each individual, calculated using the second (DD2) and the third day (DD3) in constant darkness. Phase of knock-out flies was compared with both parental controls. Colored lines represent the median phase of each group, while the length of the vector represents the standard deviation of the corresponding group (i.e., the phase coherence) (see also Tab S2). (Levene’s Test: F(4,352)= 23.59, *p*<.0001; Bartlett’s Test: *χ^2^*= 251.71, df= 4, *p*<.0001). Shapiro Normality test; F-test and Mood test (Tab S4, S5). Note: * p < 0.05, ** p < 0.01, *** p < 0.001, **** p < 0.0001.

### HSP83 depletion impairs *Pdf* transcription and PDF accumulation in s-LNv neurons

Because *Hsp83* depletion restricted to the PDF-expressing subset of clock neurons is sufficient to induce behavioural variability, we investigated if PDF expression is affected by HSP83. To analyze potential effects on *Pdf* transcription, we applied the transcriptional *Pdf-Red* reporter in which a 0.6 kb *Pdf* promoter drives the expression of the monomeric and soluble red fluorescence protein mRFP1 (Ruben et al., 2012). Although *Pdf* mRNA levels do not oscillate across the day, we dissected flies at ZT2 and quantified RFP intensity in the soma of the l-LNv, the s-LNv, and in the dorsal s-LNv projections. Homozygous *Hsp83^08445^* mutant brains showed the typical arborization pattern of the PDF-expressing LNv in the dorsal brain and the optic lobes, indicating that HSP83 is not required for the development of these neurons and circuit formation (Fig 5A, see also Tab S3). Compared to wild type and heterozygous mutant control flies, homozygous *Hsp83^08445^* mutant flies exhibited reduced RFP signals in the s- and l-LNv soma, as well as in the dorsal s-LNv projections, indicating that *Hsp83* depletion results in reduced transcription of the *Pdf* gene (Fig 5B-E, S1A). To determine effects of HSP83 depletion on PDF levels, we dissected brains of flies with *Hsp83* knock-out in all clock neurons (*Clk856 > Cas9, hsp83 sgRNA*). Anti-PDF stainings revealed that *Hsp83* disruption in the LNv did not interfere with the normal development of these clock neurons, nor with the s-LNv projections into the dorsal brain (Tab S3). Dorsal s-LNv projections of wild type flies show higher PDF levels in the early morning compared to the early evening (Park et al., 2000) (Fig 5G, S1B). Interestingly, in *Hsp83* knock-out flies this distribution was maintained and the amplitude of PDF oscillations was increased due to lower trough levels at ZT14 (Fig 5J, S1B). PDF levels in the s-LNv soma of wild type flies peak twice, once during the day (∼ZT2-ZT8) and a second time during the night (∼ZT14-ZT22), whereby the peak levels at night are higher compared to those during the day (Park et al., 2000) (Fig. 5F, S1B). Interestingly, in *Hsp83* knock-out flies, peak PDF levels occur in the morning, and trough levels are observed at ZT14 (Fig 5I, S1B).

**Fig. 5.**
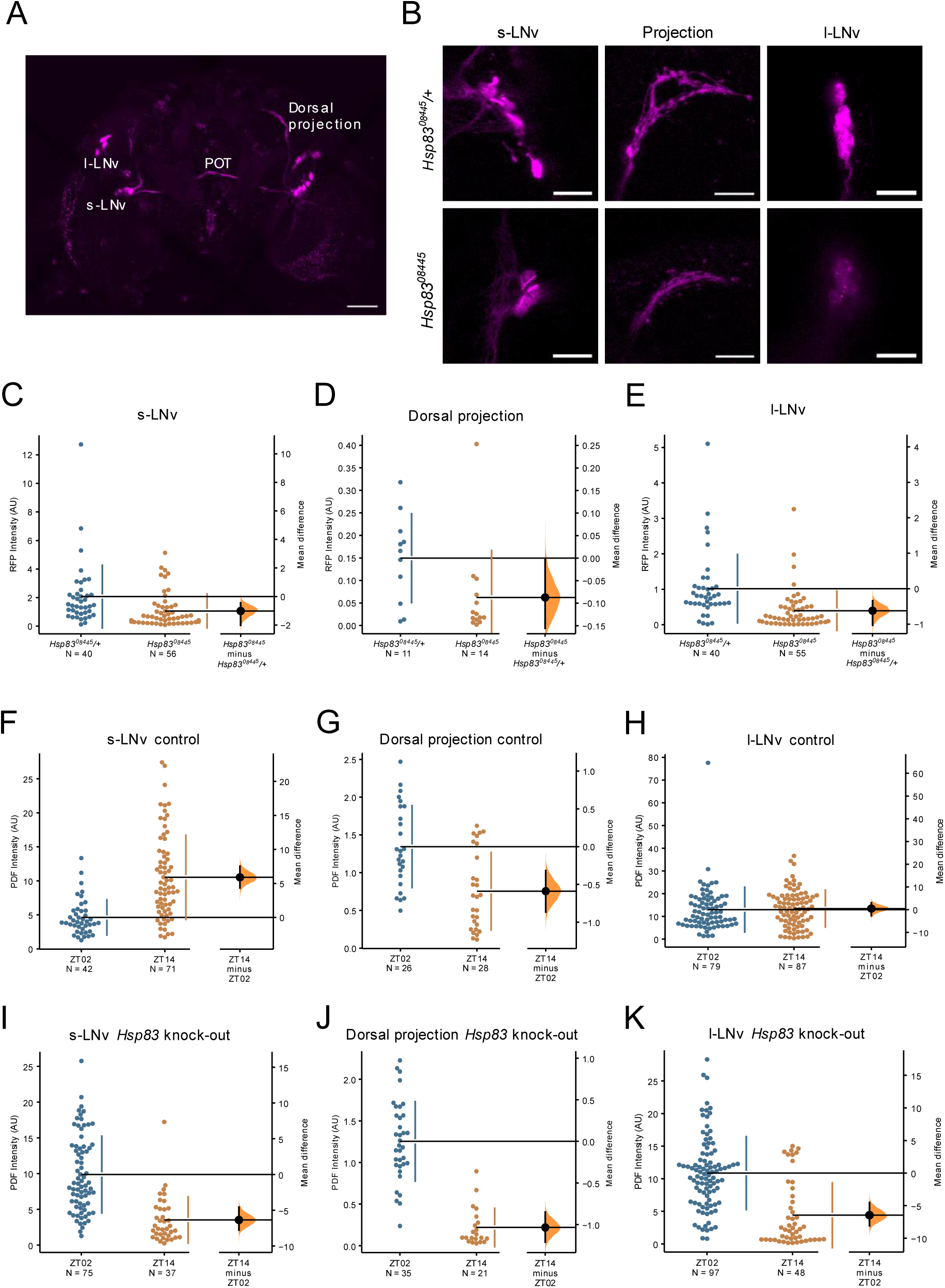
*Hsp83* regulates *Pdf* transcription and PDF levels. A) Whole brain image of a *Pdf-Red; Hsp83^08455^/Hsp83^08455^* fly, showing the s-LNv, l-LNv and the dorsal and contralateral s-LNv and l-LNv projections, respectively. POT: Posterior Optic Tract. Scale bar: 50µm. B) Representative images of the s-LNv soma, their dorsal projections and the l-LNv soma of the heterozygous and homozygous mutant *Hsp83^08445^* flies. Scale bar: 20µm. C-E) Quantification and Estimation Statistics (ES) of the RFP levels for heterozygous and homozygous mutant *Hsp83^08445^* flies for the s-LNv (C), dorsal projections (D) and l-LNv (E) (see Fig S1A for a comparison to wild type flies). F-H) Quantification and ES of PDF expression in the control *UAS-Hsp83-sgRNA/+* flies in the somas of the s-LNv (F), dorsal projections (G), and l-LNv (H). I-K) Quantification and ES of PDF expression in *Hsp83* knock-out flies (*Clk856>Cas9, Hsp83-sgRNAi*) in the s-LNv (I), dorsal projections (J) and l-LNv (K). n-numbers are indicated below each genotype. For the somas, data from single neurons is represented, and for the projections, n refers to hemispheres. In each of the panels (C-K), the left plot represents all the datapoints, with the mean (size of gap between vertical lines) and the standard deviation (length of the vertical bars) for each condition to the right. The right plot shows the mean difference of the data from the two conditions as a bootstrap 95% confidence interval (see Materials and Methods for details).

Considering that PDF release from the dorsal projections occurs during the daytime, our results indicate that in *Hsp83* knock-out flies--after its release--PDF is not restored to normal levels during the night, presumably due to reduced *Pdf* transcription (Fig 5B-E, S1B). Consistent with this interpretation, after *Hsp83* depletion we also observe lower PDF levels at ZT14 in the l-LNv soma compared to controls, which show equally high PDF levels during the day and night (Park et al., 2000) (Fig 5H, K, S1B). Finally, VRI is crucial for PDF accumulation in the dorsal s-LNv projections, and this regulation of PDF levels occurs at the posttranscriptional level (Gunawardhana and Hardin, 2017). To rule out that the reduced PDF levels we observed at ZT14 are due to altered VRI levels in *Hsp83* mutants, we performed anti-VRI stainings in the LNv of *Hsp83^08445^* and control brains at ZT20 when VRI expression is high in wild type flies (Gunawardhana and Hardin, 2017) (Fig. S1C). We observed no differences in VRI intensity between mutant and wild type LNv, further supporting a role for *Hsp83* in the transcriptional regulation of *Pdf* (Fig S1C).

### *Hsp83* mutants affect PERIOD expression rhythms in PDF-negative clock neurons

In DD, high-amplitude PER oscillations in the 5^th^ sLNv and the LNd do not depend on PDF (Yoshii et al., 2009). In fact, in *Pdf^01^* mutants PER robustly oscillates in phase in all six LNd, while in wild type PER cycles in opposite phase in the three CRY^+^ compared to the three CRY^-^ LNd (Yoshii et al., 2009). These effects of PDF-absence are nicely reflected by the expression of the *period-luciferase* reporter *8.0-luc*: This promoter-less PER encoding transgene contains intronic *per* regulatory sequences, directing PER expression to the three CRY^-^ LNd, the 5^th^-LNv, and the majority of the DN neurons (DN1-DN3) (Veleri et al., 2003; Yoshii et al., 2009). Strikingly, in a *Pdf^01^* mutant background *8.0-luc* expression in the DN groups is strongly reduced, while strong expression is observed in all six LNd and the 5^th^ s-LNv (Yoshii et al., 2009). This also explains why in *Pdf^01^* the *8.0-luc* bioluminescence rhythms are more robust and of higher amplitude compared to the wild type background (Yoshii et al., 2009). Due to the strong effects of *Hsp83* mutants on *Pdf* transcription and PDF accumulation, we analyzed *8.0-luc* expression in the *Hsp83^08445^* mutant background (Fig 6). To determine bioluminescence expression, 15 *8.0-luc* males in wild type, *Pdf^01^,* or *Hsp83^08445^* mutant background were measured for 9 days in a LumiCycle luminometer in DD at 25°C with high temporal resolution (4 min intervals, (Johnstone et al., 2022)). Strikingly, we observed a strong increase in the amplitude of *8.0-luc* oscillations in *Hsp83^08445^* mutant flies, even surpassing the effects of *Pdf^01^*(Fig 6). As evident from the raw data plotted in Fig 6A, high-amplitude oscillations continue throughout the experiment in the background of both *Pdf^01^* and *Hsp83^08445^* mutant flies, consistent with the epistatic effects of *Hsp83* on *Pdf* expression (Fig 5). Similar to *Pdf^01^*, *Hsp83^08445^* also resulted in short-period *8.0-luc* oscillations, reducing τ to 23.6 h--1.5 h shorter compared to the wild type background (Fig 6B-C, Yoshii et al 2009). In summary, these results further support that *Hsp83* positively regulates *Pdf* expression thereby explaining the molecular and behavioral phenotypes of *Hsp83* mutations.

**Fig. 6.**
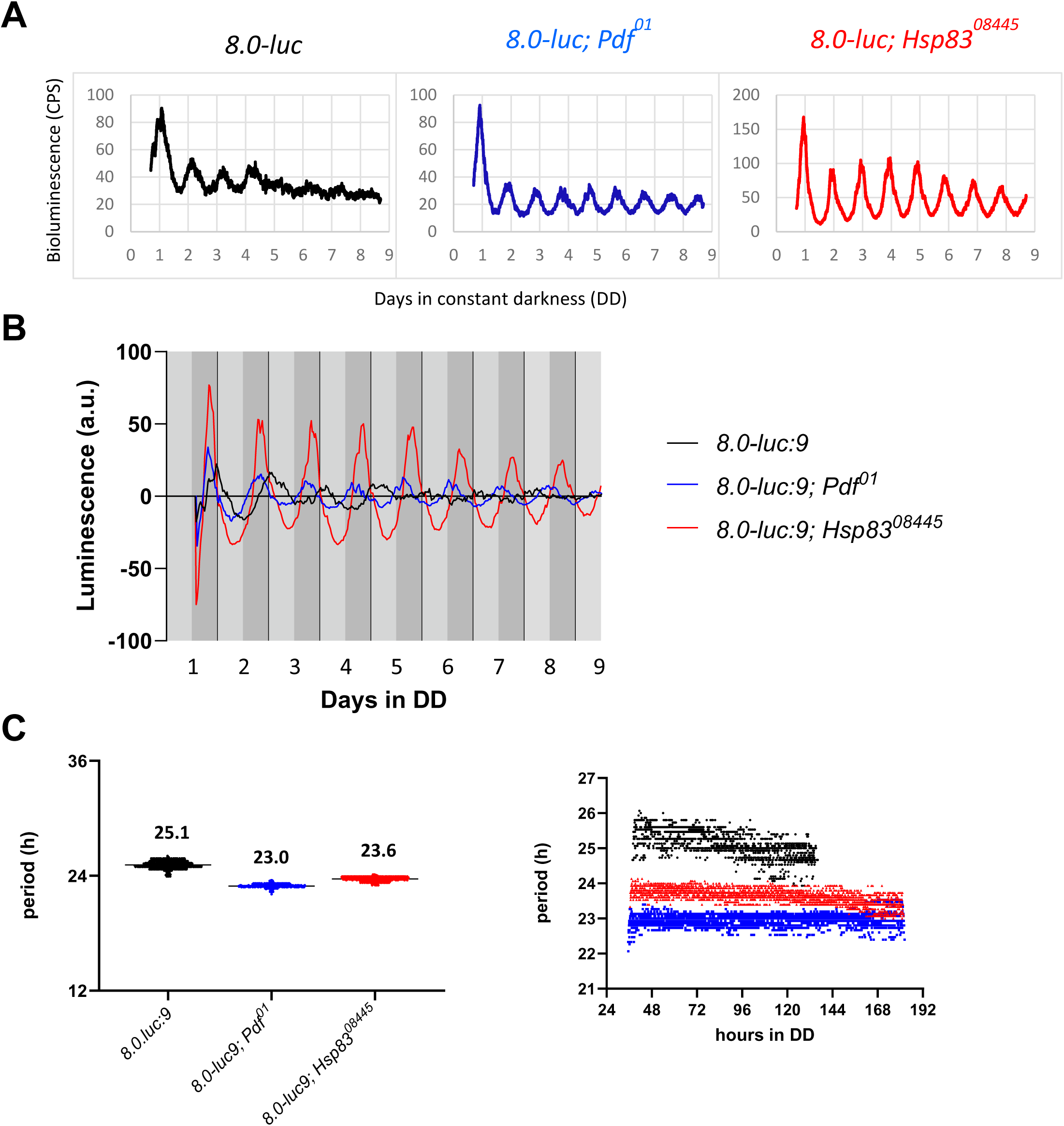
Hsp83 depletion affects PERIOD-LUCIFERASE in subsets of clock neurons. Bioluminescence recordings from *8.0-luc* flies in the genetic background indicated. 15 flies were measured for 9 days in DD at 25°C with 4 min time resolution. **A)** Raw bioluminescence data (counts per second, CPS). **B)** Detrended data (arbitrary units, a.u.). **C)** Period estimations using wavelet analysis averaged over 9 days (left) and over time (right). See text and Materials and Methods for details.

## Discussion

We show here that the chaperon protein HSP83 buffers inter-individual behavioural variability. In contrast to a previous study (Hung et al., 2009), we did not observe significant effects of hypomorphic *Hsp83* mutants on the strength of free-running behavioural rhythms, with the exception of *Hsp83* depletion in all clock neurons (Tab 1). While we have no explanation for this discrepancy, we were able to reproduce and refine the overall increased circadian plasticity. Importantly, we observed a significant decrease in phase coherence, both in *Hsp83* mutants and after *Hsp83* depletion in all clock neurons, or the PDF expressing subset only. Our results therefore pinpoint the clock neurons as cellular substrate for HSP83 function in buffering behavioural plasticity. For morphological traits, HSP83 depletion is thought to increase variation via the release of CGV, or through epigenetic mechanisms, although the underlying molecular mechanisms are not understood (Rohner et al., 2013; Rutherford and Lindquist, 1998; Sollars et al., 2003). An epigenetic mechanism is usually characterized by a maternal effect phenotypically expressed in the heterozygous offspring (Sollars et al., 2003). Because we did not observe an increase of behavioural plasticity in heterozygous *Hsp83^08445^/*+ flies resulting from a cross of homozygous *Hsp83^08445^/Hsp83^08445^* females to wild type (*iso31*) males (Fig 2, Tab 1,2), we favour the idea that Hsp83 depletion releases CGV.

We show that HSP83 is intricately linked to the regulation of PDF, a neuropeptide required for normal locomotor behaviour of fruit flies (Renn et al., 1999). *Hsp83* depletion leads to reduced *Pdf* expression at the transcriptional level, presumably resulting in slower replenishment of PDF levels after PDF release in the s-LNv terminals in the dorsal brain. While the effects on PDF expression supply a reasonable explanation for the observed plasticity, the behavioural phenotypes of *Hsp83* depletion do not match those of *Pdf^01^* mutants. Flies lacking PDF in the s-LNv display short-period behavioural rhythms in DD, which after a few days deteriorate to arrhythmicity. Moreover, PDF is required for morning anticipation (i.e., the behavioural activity increase before the lights come on in the morning), as well as for proper timing of the evening activity peak during LD (Renn et al., 1999; Shafer and Taghert, 2009). We think that altered PDF-expression in *Hsp83* mutants, as opposed to lack of PDF in *Pdf^01^* mutants are the reason for this discrepancy. PDF exerts its function after release from dense core vesicles and binding to its receptor (PDFR), which is expressed in non-PDF clock neurons, including the three CRY-positive LNd and the 5^th^ s-LNv, the DN1a, six Cry-positive DN1p, a few DN3, and the PDF-expressing s-LNv themselves (Im and Taghert, 2010). While PDF is required for synchronous and high-amplitude oscillations in most of the clock neurons (e.g., the s-LNv), other groups, like the six LNd and the 5^th^ s-LNv exhibit increased synchrony of clock gene expression and cycle with high-amplitude and short period in the absence of PDF (Lin et al., 2004; Yoshii et al., 2009). These effects of *Pdf^01^* on clock gene expression are accurately and conveniently reported by the *8.0-luc period-luciferase* reporter whose expression is largely restricted to the PDF-negative PDFR clock neurons and the Cry-negative LNd (Veleri et al., 2003; Yoshii et al., 2009) (Fig 6). We observed very similar effects of the *Hsp83^08445^* mutant on *8.0-luc* expression, including high-amplitude and short-period rhythms, further supporting HSP83 influence on PDF-signaling (Fig 6).

Our results are reminiscent of a recent study showing that transcriptional changes in *Pdf* expression contribute to circadian plasticity between different Drosophila species (Shahandeh et al., 2024). The equatorial species *D. sechellia* has lost the ability to adjust its activity to long photoperiods, which is causally linked to reduced *Pdf* expression caused by altered 5’-regulatory sequences compared to *D. melanogaster* (Shahandeh et al., 2024). Regulation of *Pdf* expression therefore appears to be an endpoint of various factors that influence circadian plasticity at the species (e.g., evolution of regulatory sequences) and individual (regulation by HSP83) level. Taken together our results provide compelling evidence for a role of HSP83 in regulating PDF expression. Given the more substantial behavioural effects of HSP83 depletion in all clock neurons compared to PDF neurons only (Tab1, Figs 3, 4), it seems likely that HSP83 has other targets, either directly within the core molecular clock mechanism, or affecting other circadian neuropeptides. For example, in the mammalian clock system HSP90 stabilizes the CYC homologue BMAL1, and fly clock neurons express at least eleven different neuropeptides in addition to PDF (Reinhard et al., 2024; Schneider et al., 2014). Identifying PDF-misregulation as consequence of *Hsp83* depletion and cause of behavioural variability, our findings set the stage for addressing the underlying molecular mechanism of HSP83 behavioural capacitor function. Our results provide evidence for a role of HSP83 in the transcriptional regulation of *Pdf*. It is known that *Pdf* transcription is indirectly regulated by the transcription factor CLK, which activates the transcription of the genes *stripe* (*sr*) and *Hormone receptor-like in 38* (*Hr38*) within the s-LNv, both of which encode Zinc Finger transcription factors (Mezan et al., 2016). Because both, SR and HR38 repress *Pdf* transcription (Mezan et al., 2016) it is unlikely that these transcription factors (and CLK) are targets of HSP83 chaperon function. Transcription factors that positively regulate *Pdf* transcription have not been identified yet, but are possibly regulated by *Hsp83* to increase behavioural plasticity under environmental stressful conditions.

## Materials and Methods

### Fly strains and maintenance

Flies were raised on fly food containing 0.7% agar, 1.0% soya flour, 8.0% polenta/maize, 1.8% yeast, 8.0% malt extract, 4.0% molasses, 0.8% propionic acid, 2.3% nipagin) in 12 h:12 h LD cycle at 60% relative humidity at 25°C. All *Hsp83* variants were obtained from the BDSC and *w iso31* flies were used as controls (Koh et al., 2008). *Hsp83^e6D^* (BDSC 5696) is an antimorphic, homozygous lethal EMS-induced allele with a single amino-acid replacement (E317K) (van der Straten et al., 1997). *Hsp83^e6A^* (BDSC 36576) is a strong hypomorphic, homozygous lethal EMS-induced allele with the single amino-acid replacement S592F (van der Straten et al., 1997). *Hsp83^j5c2^*(BDSC 12064) is an amorphic homozygous lethal allele, induced by the insertion of the *P{lacW}* transposable element into the first non-coding *Hsp83* exon, which is part of both *Hsp83-RA* and *Hsp83-RB* transcripts. *Hsp83^08445^* (BDSC 11797) is a hypomorphic allele, induced by the insertion of the *P{PZ}* transposable element into the first non-coding exon of the *Hsp83-RB* transcript. *Hsp83^08445^* is homozygous viable (presumably because the *Hsp83-RA* is not, or only mildly affected), and produces viable offspring when combined with the other three *Hsp83* alleles (Tab. 1). The *Hsp83sgRNA* line (BDSC 80843) expresses *Hsp83* specific sgRNA ubiquitously for mutagenesis of *Hsp83* under *UAS-Cas9* (BDSC 58985) control. *tim-gal4:27* (Kaneko and Hall, 2000), *Clk856-Gal4* (Gummadova et al., 2009) and *Pdf-Gal4* (Renn et al., 1999), were used to drive *Cas9* expression in all clock cells, clock neurons, or the PDF expressing LNv subset, respectively. *8.0-luc:9* flies contain a promoter-less *period-luciferase* transgene on chromosome *2,* encoding a PER-LUC fusion protein expressed in subsets of the PDF negative clock neurons DN1-3, LNd, and 5^th^ s-LNv (Veleri et al., 2003; Yoshii et al., 2009). *8.0-luc:9* was combined with the *Pdf^01^* (Yoshii et al., 2009) and *Hsp83^08445^* allele using standard genetic crosses.

### Behavior

1 to 5 days old male flies were loaded individually into locomotor glass tubes with food (4% sucrose and 2% agar) at one end and a cotton cup at the other. The glass tubes were placed into DAM2 monitors of the Drosophila Activity Monitor (DAM) system (TriKinetics), to measure locomotor activity of the flies. DAM monitors were housed in programmable environmental incubators (Percival Scientific, USA). Flies were synchronized to 12 h: 12h light-dark (LD) cycle for at least 3 days and subsequently kept in constant darkness (DD) for at least 5 more days at 25°C. As controls for each genotype, mutants, Gal4 driver, and UAS lines were crossed *w iso31*, and heterozygous offspring was analyzed.

### Behavioral analysis

We analyzed activity in 3 days in LD and 5 days in DD. Actograms were computed using Rethomics in R (Geissmann et al., 2019), while phase graphs were created using a custom-made script in R (v4.1.2; R Core Team 2021). Period, percentage of rhythmic flies, rhythmic strength (RS) and phase analysis were computed using Flytoolbox in Matlab (Levine et al., 2002). Entrainment index (EI) was computed in R (v4.1.2; R Core Team 2021), based on established protocols and correcting for the startle responses to light (Gentile et al., 2013). To calculate if flies concentrate most of their activity in anticipation of the light-dark transitions in the morning and evening, we summed up the activity during the 6 h before the respective transition, divided by the total activity 6 h before and the 6 h after each transition. In order to quantify the percentage of entrained individuals, we summed the entrained individuals who concentrate more than three quarters of their activity in the 6 hours preceding the light transitions, therefore who had an EI higher than 0.75.

To gain a better understanding of the inter-individual variability observed in our experiments, we used hidden Markov models (HMMs) analysis to infer activity states from light barrier crossings (Feldmann et al., 2023). We modelled data from the mutants and cell-specific knock-out separately, using 2-state HMMs to distinguish active and inactive behaviour. The (negative binomial) state-dependent distributions of locomotor activity are estimated separately for LD and DD, and were determined based on the controls only. Subsequently, we fixed the distributional parameters for fitting HMMs to the other genotypes to ensure comparability of the activity states in LD and DD between genotypes. To investigate temporal activity patterns, we modelled the state-switching probabilities as a function of time of day for each genotype, including random intercepts to account for individual differences. For reasons of parsimony, we restricted the random effects variances to be the same for the two state-switching probabilities. All models were implemented using functions from the R package LaMa (Koslik, 2025). Based on the fitted HMMs, we derived the model-implied percentage of time spent in each state as a function of the time of day (Koslik et al., 2023)and calculated its variance between individuals at the morning and evening peaks (determined based on the control genotype *iso31*) for each genotype in LD and DD, respectively.

### Bioluminescence assay

D-luciferin potassium salt (Biosynth) was mixed with standard fly food to a final concentration of 15 mM in Drosophila culture plates (Actimetrics). Luminescence of 15 males per plate was measured every 4 min for 7-8 days in DD at 25°C with a the LumiCycle 32 Color (Actimetrics). Actimetrics analysis software was used to normalize the exponential decay, data were exported into .csv files and custom python code was used to organize luminescence data into 30-minute bins (LABLv9.py; www.top-lab.org/downloads), and to quantify periods of oscillations using a Morlet wavelet fit (waveletsv4.py; www.top-lab.org/downloads). Data were plotted using Graphpad Prism 10. For details see (Johnstone et al., 2022).

### Immunohistochemistry

For the staining of the cell-specific Hsp83 knock-out, we first entrained flies for 5 days in 12 hours light 12 hours dark (LD). After that, we fixed flies at ZT02and ZT14. Flies were fixed in 4% PFA for 2.5 h at room temperature (RT). After fixation, the samples were washed 6 times with 0.1 M phosphate buffer (pH 7.4) with 0.1% Triton X-100 (PBS-T) at RT. Brains were dissected in PBS, were then blocked with 5% goat serum in 0.1% PBS-T for 2 h at RT and stained with mouse anti-PDF C7 (DSHB, 1:500) or guinea pig anti-VRI (1:2000) (Glossop et al., 2003) 5% goat serum and 0.5% PBST for at least 48 h at 4⁰C. After washing 3 times in PBS-T, the samples were incubated at 4°C overnight with goat anti-mouse AlexaFluor 488 nm or anti-guinea pig Alexa Fluor 568 (both 1:600) in PBS-T. Brains were washed 3 times in PBS-T before being mounted in Vectashield. Brains of 1 to 2 days old *Pdf-Red* flies were dissected in a physiological medium (Buhl et al., 2016), mounted in ProLong Gold (Thermo Fisher Scientific), and imaged directly. All images were taken using a Leica SP8 confocal microscope, keeping the same settings for each experiment.

### Immunostaining quantifications

For quantification of RFP and PDF intensity, pixel intensity of mean and background staining in each neuronal group was measured by ImageJ, FIJI (Schindelin et al., 2012). Soma intensity was quantified drawing one vector in the center of the neuron and then we used the same vector to measure the corresponding background intensity. For PDF projections, we extracted a stack of the 10 Z slides where the s-LNv dorsal projections are showed at their maximum intensity. Then, we drew a vector at the center of the projection and used the same vector to measure the background intensity. Data were normalized by dividing the values by the length of the vector and consequently by subtracting the background signal. Quantification of VRI intensity was performed as described in (Ogueta et al., 2018). In brief, for each cell three measurements were taken, as well as 3 measurements of the background for the corresponding slice for background subtraction.

### Statistical analysis

The statistical tests used are listed in each figure and are summarized in supplementary tables (Tab S4-S6). Statistics for phase analysis (Fig 2, 4) were computed in R (R Core Team, 2021). We used the Shapiro-Wallis test to check normality (Table S4). When comparing phase variabilities, we applied different statistical tests for variance: Levene’s Test for Homogeneity of Variance, Bartlett’s Test of Homogeneity of Variance (both reported in the figure label), the F-test of Equality of Variances when the data were normally distributed, or the non-parametric Mood Two-Sample Test of Scale whenever the data were not normally distributed (both reported in Table S5). For staining intensity, we used estimation statistics (Ho et al., 2019) (Tab S6).

## Acknowledgements

We thank Paul Hardin for anti-VRI antibody and Justin Blau for Pdf-Red flies. We thank Reshma R, Tobias Prüser and Joachim Kurtz for discussions and for sharing unpublished results. This work was funded by the German Research Foundation (DFG) as part of the SFB TRR 212 (NC3, Project number 316099922)—Project numbers 471672756 and 396782756 to Ralf Stanewsky and Roland Langrock, respectively).

