## Supplementary Information for "Hsp90 buffers behavioral plasticity by regulating *Pdf* transcription in clock neurons of *Drosophila melanogaster*"

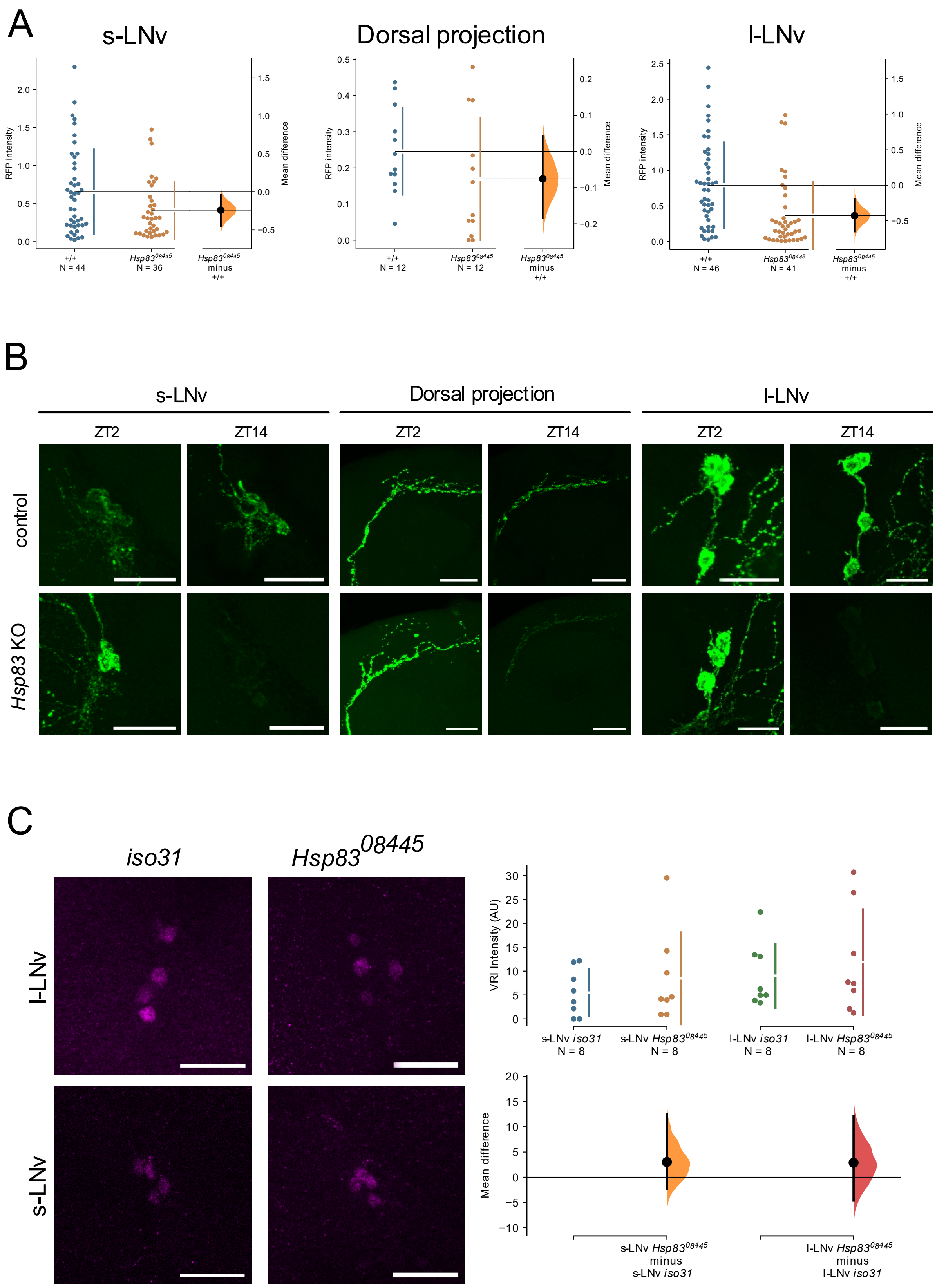


**Supplementary Figure 1**: A) Quantification of the RFP levels in *Pdf-Red; +/+* compared to *Pdf-red; Hsp83*$^{08445}$/*Hsp83*$^{08445}$ flies for the s-LNv, dorsal projection and l-LNv, using Estimation Statistics. B) Representative images of anti-PDF staining of the s-LNv, dorsal projections, and l-LNv in *Hsp83* knock out (*Clk856 > Cas9, hsp83 sgRNA*) and control flies (*hsp83 sgRNA/+*). Scale bar: 20µm. C) Left:

Representative images of anti-VRI staining in the l-LNv and s-LNv of *iso31* and $Hsp^{83084457}/Hsp^{8308445}$ flies. Scale bar: 20µm. Right: Quantification of the intensity of VRI staining in the s-LNv and l-LNv in 8 hemispheres, using Estimation Statistics.

**Table S1. Viability of *Hsp83* allelic combinations.**

| | | Males | | | | |
|---|---|---|---|---|---|---|
| | | *Df(3L)BSC672* | *Hsp83*$^{08445}$ | *Hsp83*$^{e6A}$ | *Hsp83*$^{e6D}$ | *Hsp83*$^{j5c2}$ |
| Females | *iso31* | + | + | + | + | + |
| | *Hsp83*$^{08445}$ | + | + | + | + | + |
| | *Hsp83*$^{e6A}$ | - | + | - | - | - |
| | *Hsp83*$^{e6D}$ | - | + | - | - | - |
| | *Hsp83*$^{j5c2}$ | - | + | - | - | - |
| | *Df(3L)BSC672* | - | + | - | - | - |

| | | Females | | | | |
|---|---|---|---|---|---|---|
| | | *Df(3L)BSC672* | *Hsp83*$^{08445}$ | *Hsp83*$^{e6A}$ | *Hsp83*$^{e6D}$ | *Hsp83*$^{j5c2}$ |
| Males | *iso31* | + | + | + | + | + |
| | *Hsp83*$^{08445}$ | + | + | + | + | + |
| | *Hsp83*$^{e6A}$ | - | + | - | - | - |
| | *Hsp83*$^{e6D}$ | - | + | - | - | - |
| | *Hsp83*$^{j5c2}$ | - | + | - | - | - |
| | *Df(3L)BSC672* | - | + | - | - | - |

| | | Males | | | | |
|---|---|---|---|---|---|---|
| | | *iso31* | *Hsp83 sgRNA* | *TimGal4:27; UASCas9* | *Clk856Gal4; UASCas9* | *PdfGal4; UASCas9* |
| Females | *iso31* | + | + | + | + | + |
| | *Hsp83 sgRNA* | + | + | - | + | + |

| | | Females | | | | |
|---|---|---|---|---|---|---|
| | | *iso31* | *Hsp83 sgRNA* | *TimGal4:27; UASCas9* | *Clk856Gal4; UASCas9* | *PdfGal4; UASCas9* |
| Males | *iso31* | + | + | + | + | + |
| | *Hsp83 sgRNA* | + | + | - | + | + |

*Note:* Viability is indicated with a + and crosses producing no transheterozygous mutant offspring are indicated with -.

**Table S2. Circular Phase plot and HMM analysis reveal increased variability of evening activity phase after HSP83 depletion in constant darkness (DD)**

| Genotype | Phase in circadian time (mean± SD) | SD of time spent in the active state |
|---|---|---|
| *iso31* | $10.3 \pm 0.73$ | 0.02 |
| *Hsp83$^{08445}$/+* | $10.9 \pm 0.73$ | 0.01 |
| *Hsp83$^{08445}$/Hsp83$^{08445}$* | $9.6 \pm 1.11$ | 0.19 |
| *Hsp83$^{e6A}$/+* | $10.9 \pm 0.64$ | 0.03 |
| *Hsp83$^{e6A}$/Hsp83$^{08445}$* | $9.1 \pm 2.40$ | 0.25 |
| *Hsp83$^{e6D}$/+* | $11.8 \pm 1.01$ | 0.08 |
| *Hsp83$^{e6D}$/Hsp83$^{08445}$* | $11.8 \pm 1.37$ | 0.19 |
| *Hsp83$^{j5c2}$/+* | $9.2 \pm 1.02$ | 0.03 |
| *Hsp83$^{j5c2}$/Hsp83$^{08445}$* | $8.8 \pm 1.57$ | 0.06 |
| *Hsp83 sgRNA/+;* | $10.3 \pm 0.97$ | 0.06 |
| *Clk856-Gal4>UAS-Cas9,+* | $9.8 \pm 0.98$ | 0.01 |
| *Clk856-Gal4>UAS-Cas9,Hsp83 sgRNA* | $9.9 \pm 3.83$ | 0.25 |
| *Pdf-Gal4>UAS-Cas9,+* | $9.9 \pm 0.74$ | 0.01 |
| *Pdf-Gal4>UAS-Cas9,Hsp83 sgRNA* | $10.9 \pm 2.58$ | 0.12 |

Notes: *SD of model-implied percentage of time spent in the active state were quantified based on the HMM analysis.*

**Tab S3. Number of PDF projections, s-LNvs and l-LNvs neurons per hemisphere in cell-specific *Hsp83* knock-out and parental control.**

| Neurons | Genotype | ZT | # hemisphere | # neurons per hemisphere (mean±SD) | % Hemispheres with abnormalities (n. abnormal projections/total hemipheres) |
|---|---|---|---|---|---|
| s-LNv | *Hsp83 sgRNA/+* | 02 | 17 | $2.5 \pm 0.72$ | - |
| | | 14 | 21 | $3.4 \pm 0.74$ | - |
| | *Clk856-Gal4>UAS-Cas9, Hsp83 sgRNA* | 02 | 28 | $2.7 \pm 1.12$ | - |
| | | 14 | 14 | $2.6 \pm 1.01$ | - |
| l-LNv | *Hsp83 sgRNA/+* | 02 | 22 | $3.6 \pm 0.50$ | - |
| | | 14 | 24 | $3.6 \pm 0.49$ | - |
| | *Clk856-Gal4>UAS-Cas9, Hsp83 sgRNA* | 02 | 32 | $3.0 \pm 0.97$ | - |
| | | 14 | 17 | $2.8 \pm 0.88$ | - |
| PDF dorsal projections | *Hsp83 sgRNA/+* | - | 56 | - | 0.04% (2/56) |
| | *Clk856-Gal4>UAS-Cas9, Hsp83 sgRNA* | - | 53 | - | 0.04% (2/53) |

**Table S4. Shapiro-Wilk Normality Test.**

| Parameters | Statistic (W) | p-value | Signif. | Fig. |
|---|---|---|---|---|
| iso31 | 0.98 | 0.24 | ns | 2 |
| *Hsp83*$^{08445}$*/+* | 0.98 | 0.21 | ns | |
| *Hsp83*$^{08445}$*/ Hsp83*$^{08445}$ | 0.95 | < 0.01 | ** | |
| *Hsp83*$^{e6A}$*/+* | 0.91 | 0.001 | * | |
| *Hsp83*$^{e6A}$*/ Hsp83*$^{08445}$ | 0.96 | 0.35 | ns | |
| *Hsp83*$^{e6D}$*/+* | 0.91 | < 0.001 | *** | |
| *Hsp83*$^{e6D}$*/ Hsp83*$^{08445}$ | 0.88 | < 0.0001 | **** | |
| *Hsp83*$^{j5c2}$*/+* | 0.98 | 0.42 | ns | |
| *Hsp83*$^{j5c2}$*/ Hsp83*$^{08445}$ | 0.77 | < 0.0001 | **** | |
| *Hsp83 sgRNA/+* | 0.92 | < 0.0001 | **** | 4 |
| *Clk856 Gal4> UAS Cas9, +* | 0.98 | 0.37 | ns | |
| *Clk856 Gal4 > UAS Cas9, Hsp83 sgRNA* | 0.92 | 0.02 | * | |
| *Pdf Gal4 > UAS Cas9, +* | 0.97 | 0.08 | ns | |
| *Pdf Gal4 > UAS Cas9, Hsp83 sgRNA* | 0.90 | < 0.0001 | **** | |

**** $p < .0001$, *** $p < .001$, ** $p < .01$, * $p < .05$, ns $p > .05$.

**Table S5. F test to compare two variances and Mood Two-Sample Test of Scale.**

| Group 1 | Group 2 | F Test | Mood test (Z) | p-value | Signif. | Fig. |
|---|---|---|---|---|---|---|
| iso31 | *Hsp83*$^{08445}$*/+* | $F_{[84, 68]}$=0.98 | - | 0.92 | ns | 2 |
| | *Hsp83*$^{08445}$*/ Hsp83*$^{08445}$ | - | 2.17 | 0.03 | * | |
| | *Hsp83*$^{e6A}$*/+* | - | -0.92 | 0.36 | ns | |
| | *Hsp83*$^{e6A}$*/ Hsp83*$^{08445}$ | $F_{[24,68]}$=10.64 | - | < 0.0001 | **** | |
| | *Hsp83*$^{e6D}$*/+* | - | 2.42 | 0.01 | * | |
| | *Hsp83*$^{e6D}$*/ Hsp83*$^{08445}$ | - | 3.06 | <0.01 | ** | |
| | *Hsp83*$^{j5c2}$*/+* | $F_{[49, 68]}$=1.92 | - | 0.01 | * | |
| | *Hsp83*$^{j5c2}$*/ Hsp83*$^{08445}$ | - | 1.09 | 0.27 | ns | |
| *Hsp83*$^{08445}$*/+* | *Hsp83*$^{08445}$*/ Hsp83*$^{08445}$ | - | 0.85 | 0.40 | ns | |
| | *Hsp83*$^{e6A}$*/ Hsp83*$^{08445}$ | $F_{[24,84]}$=10.87 | - | < 0.0001 | **** | |
| | *Hsp83*$^{e6D}$*/ Hsp83*$^{08445}$ | - | 3.99 | < 0.0001 | **** | |
| | *Hsp83*$^{j5c2}$*/ Hsp83*$^{08445}$ | - | 2.17 | 0.03 | * | |
| *Hsp83*$^{e6A}$*/+* | *Hsp83*$^{e6A}$*/ Hsp83*$^{08445}$ | - | 3.76 | < 0.001 | *** | |
| *Hsp83*$^{e6D}$*/+* | *Hsp83*$^{e6D}$*/ Hsp83*$^{08445}$ | - | 1.14 | 0.25 | ns | |
| *Hsp83*$^{j5c2}$*/+* | *Hsp83*$^{j5c2}$*/ Hsp83*$^{08445}$ | - | 0.20 | 0.84 | ns | |
| *Hsp83sgRNA+* | *Clk856 Gal4>UAS-Cas9,+* | - | -1.06 | 0.29 | ns | 4 |
| | *Pdf Gal4>UAS-Cas9,+* | - | -2.78 | < 0.01 | ** | |
| | *Clk856 Gal4>UAS-Cas9, Hsp83 sgRNA* | - | 6.53 | < 0.0001 | **** | |
| | *Pdf Gal4>UAS-Cas9, Hsp83 sgRNA* | - | 5.83 | < 0.0001 | **** | |
| *Clk856 Gal4> UAS Cas9, +* | *Clk856 Gal4>UAS-Cas9, Hsp83 sgRNA* | - | 5.95 | < 0.0001 | **** | |
| *Pdf Gal4 > UAS Cas9, +* | *Pdf Gal4 >UAS-Cas9, Hsp83 sgRNA* | - | 6.43 | < 0.0001 | **** | |

*Note:* **** $p$ < .0001, *** $p$ < .001, ** $p$ < .01, * $p$ < .05, ns $p$ >.05.

**Table S6. Estimation statistics signal intensity.**

| Group | Genotype | ZT | Intensity (mean±SEM) | Difference intensity | 95%CI | | Fig. |
|---|---|---|---|---|---|---|---|
| | | | | | low | high | |
| s-LNv | *Hsp83*$^{08445}$ | 02 | 1.1 ± 0.2 | -1.01 | -2.01 | -0.4 | 5C |
| | *Hsp83*$^{08445}$/+ | | 2.1 ± 0.3 | | | | |
| Dorsal projection | *Hsp83*$^{08445}$ | 02 | 0.1 ± 0.03 | -0.09 | -0.2 | -0.01 | 5D |
| | *Hsp83*$^{08445}$/+ | | 0.15 ± 0.03 | | | | |
| l-LNv | *Hsp83*$^{08445}$ | 02 | 0.4 ± 0.1 | -0.6 | -1.03 | -0.3 | 5E |
| | *Hsp83*$^{08445}$/+ | | 1.02 ± 0.1 | | | | |
| s-LNv | *Hsp83sgRNA+* | 02 | 4.6 ± 0.40 | 5.9 | 4.3 | 7.5 | 5F |
| | | 14 | 10.5 ± 0.74 | | | | |
| Dorsal projection | | 02 | 1.3 ± 0.11 | - 0.6 | - 0.9 | - 0.3 | 5G |
| | | 14 | 0.7 ± 0.10 | | | | |
| l-LNv | | 02 | 13.0 ± 1.11 | 0.48 | - 2.7 | 2.9 | 5H |
| | | 14 | 13.4 ± 0.87 | | | | |
| s-LNv | *Clk856 Gal4>UAS-Cas9, Hsp83 sgRNA* | 02 | 9.9 ± 0.62 | - 6.3 | - 7.8 | - 4.6 | 5I |
| | | 14 | 3.5 ± 0.53 | | | | |
| Dorsal projection | | 02 | 1.2 ± 0.08 | - 1.03 | - 1.2 | - 0.8 | 5J |
| | | 14 | 0.2 ± 0.05 | | | | |
| l-LNv | | 02 | 10.9 ± 0.57 | - 6.5 | - 8.1 | - 4.5 | 5K |
| | | 14 | 4.4 ± 0.72 | | | | |
| s-LNv | *Hsp83*$^{08445}$ | 02 | 0.4 ± 0.1 | -0.2 | -0.5 | -0.04 | S1A |
| | *+/+* | | 0.6 ± 0.1 | | | | |
| Dorsal projection | *Hsp83*$^{08445}$ | 02 | 0.2 ± 0.05 | -0.08 | -0.2 | 0.05 | S1A |
| | *+/+* | | 0.25 ± 0.03 | | | | |
| l-LNv | *Hsp83*$^{08445}$ | 02 | 0.4 ± 0.1 | -0.4 | -0.6 | -0.2 | S1A |
| | *+/+* | | 0.8 ± 0.1 | | | | |
| s-LNv | *Hsp83*$^{08445}$ | 20 | 8.5 ± 3.4 | 3 | -2.3 | 12.4 | S1B |

| | | | | | | | |
|---|---|---|---|---|---|---|---|
| | *iso31* | | 5.5 ± 1.2 | | | | |
| l-LNv | *Hsp83*$^{08445}$ | 20 | 11.9 ± 3.9 | 2.9 | -4.6 | 12.1 | S1B |
| | *iso31* | | 9 ± 2.4 | | | | |
